# OptoDyCE-plate as an affordable high throughput imager for all optical cardiac electrophysiology

**DOI:** 10.1101/2023.08.29.555447

**Authors:** Yuli W. Heinson, Julie L. Han, Emilia Entcheva

**Affiliations:** Department of Biomedical Engineering, The George Washington University Washington, DC 20037

**Keywords:** high-throughput, all-optical electrophysiology, optogenetics, cardiotoxicity, human iPSC-CMs, drug testing, low cost, CRISPRi, cardiac tissue engineering

## Abstract

We present a simple low-cost system for comprehensive functional characterization of cardiac function under spontaneous and paced conditions, in standard 96 and 384-well plates. This full-plate actuator/imager, OptoDyCE-plate, uses optogenetic stimulation and optical readouts of voltage and calcium from all wells in parallel. The system is validated with syncytia of human induced pluripotent stem cell derived cardiomyocytes, iPSC-CMs, grown as monolayers, or in quasi-3D isotropic and anisotropic constructs using electrospun matrices, in 96 and 394-well format. Genetic modifications, e.g. interference CRISPR (CRISPRi), and nine compounds of acute and chronic action were tested, including five histone deacetylase inhibitors (HDACis). Their effects on voltage and calcium were compared across growth conditions and pacing rates. We also demonstrated deployment of optogenetic cell spheroids for point pacing to study conduction in 96-well format, and the use of temporal multiplexing to register voltage and calcium simultaneously on a single camera in this stand-alone platform. Opto-DyCE-plate showed excellent performance even in the small samples in 384-well plates, in the various configurations. Anisotropic structured constructs may provide some benefits in drug testing, although drug responses were consistent across tested configurations. Differential voltage vs. calcium responses were seen for some drugs, especially for non-traditional modulators of cardiac function, e.g. HDACi, and pacing rate was a powerful modulator of drug response, highlighting the need for comprehensive multiparametric assessment, as offered by OptoDyCE-plate. Increasing throughput and speed and reducing cost of screening can help stratify potential compounds early in the drug development process and accelerate the development of safer drugs.

## Introduction

Cardiotoxicity testing is central to drug development and has been part of preclinical assessment for all compounds in the pipeline. Recently, FDA announced that pre-clinical animal testing will not be required for moving to clinical trials, and there is intense activity to support new approach methods (NAMs) for toxicity testing and drug development [1–5]. These include high throughput (HT) in vitro assays with human induced pluripotent stem cells (iPSC-CMs), in silico methods, population studies and data mining. Human data are believed to provide more relevant information for responses to drugs and toxins compared to animal studies.

Technological developments in automation in optical sensing have advanced the methods for in vitro cardiotoxicity testing [6–8]. For example, a high-throughput imager was developed for incubator deployment based on parabolic mirror multiplexing using low cost components[8], and previously published dye-free imaging of cardiac waves [9]. Optogenetics and optical actuation[10, 11] add the possibility to control rate, beyond passive observations of spontaneous activity, thereby leading to more comprehensive all-optical electrophysiology assays [12–17]. For example, multiplexed patterned islands with various optical sensors within smaller FOV allowed multiparametric functional evaluation[18]. Overall, optical methods for cardiotoxicity testing are compatible with multicellular samples and engineered tissue, provide higher information content and may lead to better predictions compared to multielectrode arrays or other tools. Most earlier implementations of these methods have involved some serial scanning and/or multiplexing.

However, over the last 2-3 years, a surge of methodologies with increased parallelism and speed is seen, often including expansion of the field of view (FOV), in some cases, to imaging a full plate simultaneously. A sophisticated Swarm imager for all-optical electrophysiology was developed using several partitions of a HT plate [19]. Also plate partitions were used (3.5 x 2.0cm area) in cardiac electrophysiology plate reader, capturing spontaneous activity (no pacing) with genetically-encoded sensors[20]. This offered longitudinal electrophysiological measurements from 96 wells of a standard multi well plate. Using a motorized XY stage, sequential partitions were imaged, so that the recording from all 96 wells was completed in 5 minutes. A true full-plate imager was achieved using optical contractility measurements in small engineered cardiac tissues in 96 well plate via lens array mosaic projection, that allowed better focusing and better mapping of the relevant areas in the FOV [21]. All-optical electrophysiology application was reported using trans-illumination-based 24-well plate imager MULTIPLE, capturing action potential propagation under optogenetic stimulation[22]. Finally, scalable optogenetic perturbations in the incubator setting have also been pursued, e.g. by using optoplate-96 with LED matrices[23].

Our current study presents a new on-axis epifluorescence full-plate imager for all-optical electrophysiology (OptoDyCE-plate) that operates with standard 96 and 384-well plates. This high-throughput solution distinguishes itself from the recent developments with its simplicity, low cost and comprehensiveness of evaluation. The latter is demonstrated with combined voltage and calcium sensing under optogenetic pacing in glass-bottom HT plates as well as in quiasi-3D anisotropic and isotropic human iPSC-CM syncytia.

## Results

### OptoDyCE-plate system characterization

The developed OptoDyCE-plate is a full-plate imager for high-throughput functional electrophysiological testing. This on-axis epifluorescence-based optical actuator/sensor device is an extension of previous microscope-based systems we have reported[12, 13], but it is a highly parallel stand-alone version able to image while optically pacing full standard 96-well and 384-well plates. We show how to construct such HT system using off-the-shelf low-cost components (including a low-cost camera) for less than $20,000. Using spectrally compatible optical sensors and actuators, it is possible to optically image voltage and calcium under optogenetic pacing. **Figure 1** presents OptoDyCE-plate along with its characterization.

**Fig. 1.**
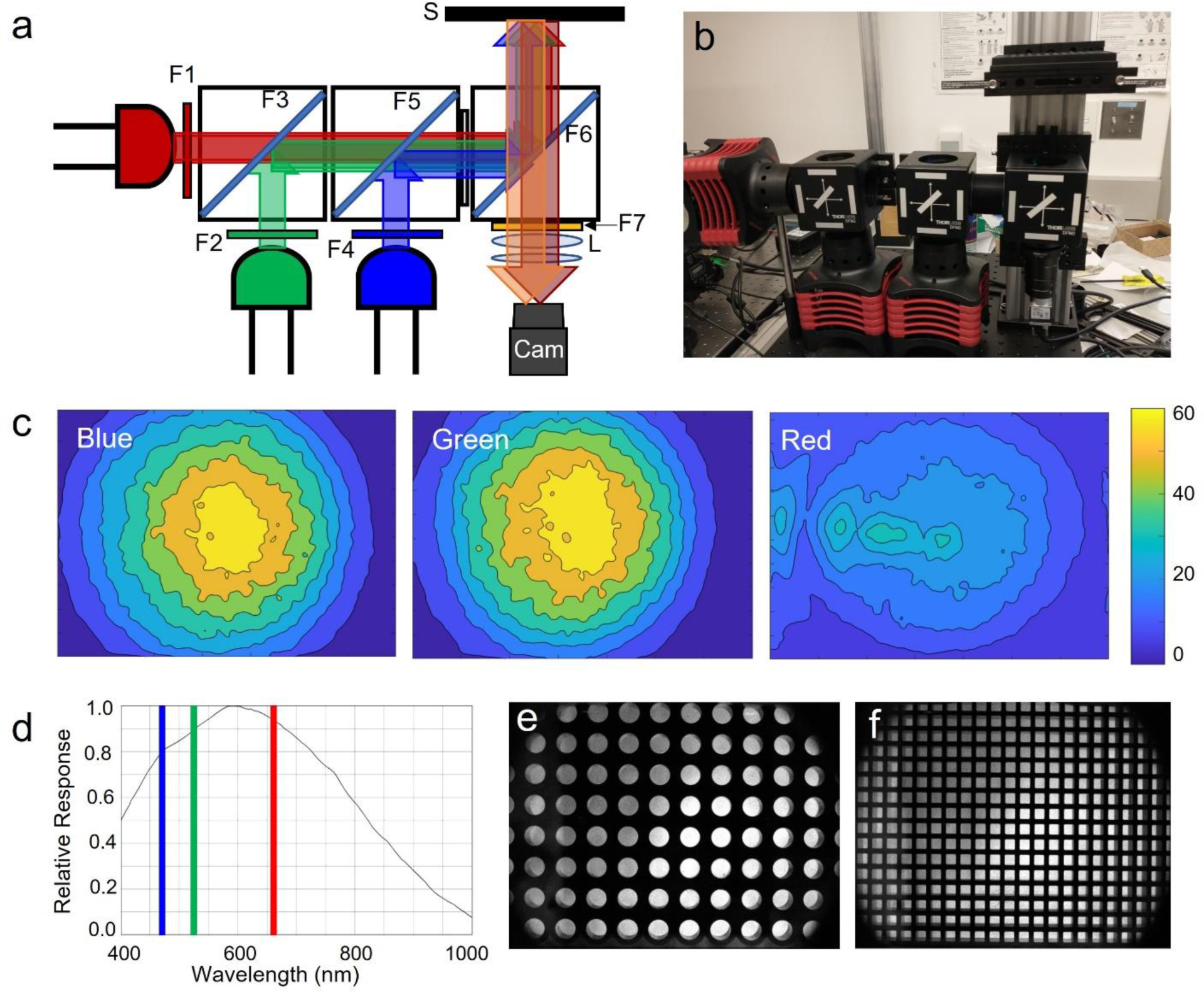
Low-cost high throughput (HT) all-optical plate imager OptoDyCE-plate. (a) Schematic diagram of the imaging system. Green LED is used for excitation of calcium-responsive optical or optogenetic sensors (Rhod4, jR-GECO), red LED for excitation of optical NIR voltage sensors, e.g. BeRST1, and blue LED is used for optogenetic stimulation. Notations include the sample stage – S, CMOS camera – Cam. Camera lens – L. Filters include: F1: 655/30; F2: 525/30; F3: 605LP dichroic; F4: 470/24; F5: 495LP dichroic; F6: 473/532/660 dichroic; F7: 595/40+700LP. (b) A photo of the imaging system structured on a 12” x 12” aluminum breadboard. Total cost was approximately $20,000, including the camera, three light sources and drivers, optical and mechanical components. (c) Irradiance profiles at the camera from the blue, green, and red LEDs, respectively, obtained by placing a piece of white printing paper on the sample stage. (d) Spectral response for the low cost Basler camera; the blue, green, and red LEDs wavelengths are indicated with the corresponding color bars. Images of a 96 well plate (e) and a 384 well plate (f) on the HT imager with room light on.

Light source irradiance was characterized across the field of view, by placing a piece of white printing paper on the sample stage and taking a still image when illuminated. **Figure 1c** shows the relative irradiance profiles seen by the camera. Due to the dichroic and emission filters (F6 and F7) in front of the camera, blocking certain wavelengths, the real irradiance at the sample position would be different, however, the relative irradiance profiles are still informative. During the irradiance characterization, the red LED was set at maximum power yielding 19 mW/cm^2^ measured with a power meter (PM100, Thorlabs). To not saturate the camera, the green and blue LEDs were set at less than their maximum powers during the irradiance characterization. The powers at the center of the sample stage from the green and blue LEDs were 11 mW/cm^2^ and 32 mW/cm^2^, respectively. During functional experiments, the red and pulsed blue LEDs were at their maximum powers; the green LED was usually set at ∼70% of the maximum power to produce good signal to noise ratio (SNR) without inducing spectral crosstalk by activating the optogenetic actuator[17]. Light irradiance limitations, especially for the red LED, impacted signal at the periphery. However, the central portion of the FOV showed excellent SNR.

A very low-cost machine vision Basler camera was used here, as in some of our recent studies[17, 24]. **Figure 1d** shows the camera’s relative spectral responses with respect to the wavelength, adapted from published specifications, with the central wavelengths of the three light sources indicated by the color bars. **Figure 1e** and **f**, present images of a 96-well plate and a 384-well plate, respectively, through the HT imager. Based on the illumination profiles from the three LEDs (**Figure 1c**), to maintain best SNR, usually 6 x 8 wells at the center of 96 well plates were recorded. For a 384-well plate, we comfortably recorded signals from at least 100 wells in parallel (10 x10 samples). Image resolution of the HT plate imager is 132µm/pixel. The limitations in resolution and FOV were based on the low-cost camera and the single illumination LED per channel. Scaling up or upgrading either can directly improve these characteristics.

Initially, we tested OptoDyCE-plate with a 96-well glass-bottom plate using the central 20 (4 x 5) wells with human induced pluripotent stem cell derived cardiomyocytes (hiPSC-CMs), genetically modified with ChR2 for optical actuation. We recorded optically both V_m_ and [Ca^2+^]_i_ activities from the 20 wells. Samples were paced optically with 0.5, 0.75, 1.0, 1.2 and 1.5Hz, while collecting voltage and calcium responses. All 20 wells followed each of the optical pacing frequencies. **Figure 2a** shows example V_m_ and [Ca^2+^]_i_ fluorescence traces for each optical pacing frequency. The traces were from the same well, indicated with squares in **Figure 2b**. We quantified the SNR for V_m_ and [Ca^2+^]_i_ traces from each well, color coded as shown in **Figure 2b**. **Figure 2c** and **d** present the SNR of V_m_ and [Ca^2+^]_i_, respectively, as a function of pacing frequency. SNRs are ∼30 for V_m_ and ∼40 [Ca^2+^]_i_. The SNR was obtained by analyzing the Fourier spectrum after subtracting the DC component:

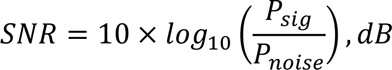

where *P_sig_* is the power at the corresponding pacing frequency and *P_noise_* is the lowest power between the first and the second harmonic. SNR variation did not have a specific pattern related to the illumination profile; it most likely depends on the precise delivery of the optical sensors, which in our case was manual. We found a statistically significant difference in SNR results by both V_m_/[Ca^2+^]_i_ labeling and by pacing frequency; the interaction between these terms was also significant, suggesting the particular optical sensor and the pacing frequency both affect the SNR of the cells imaged.

**Fig. 2.**
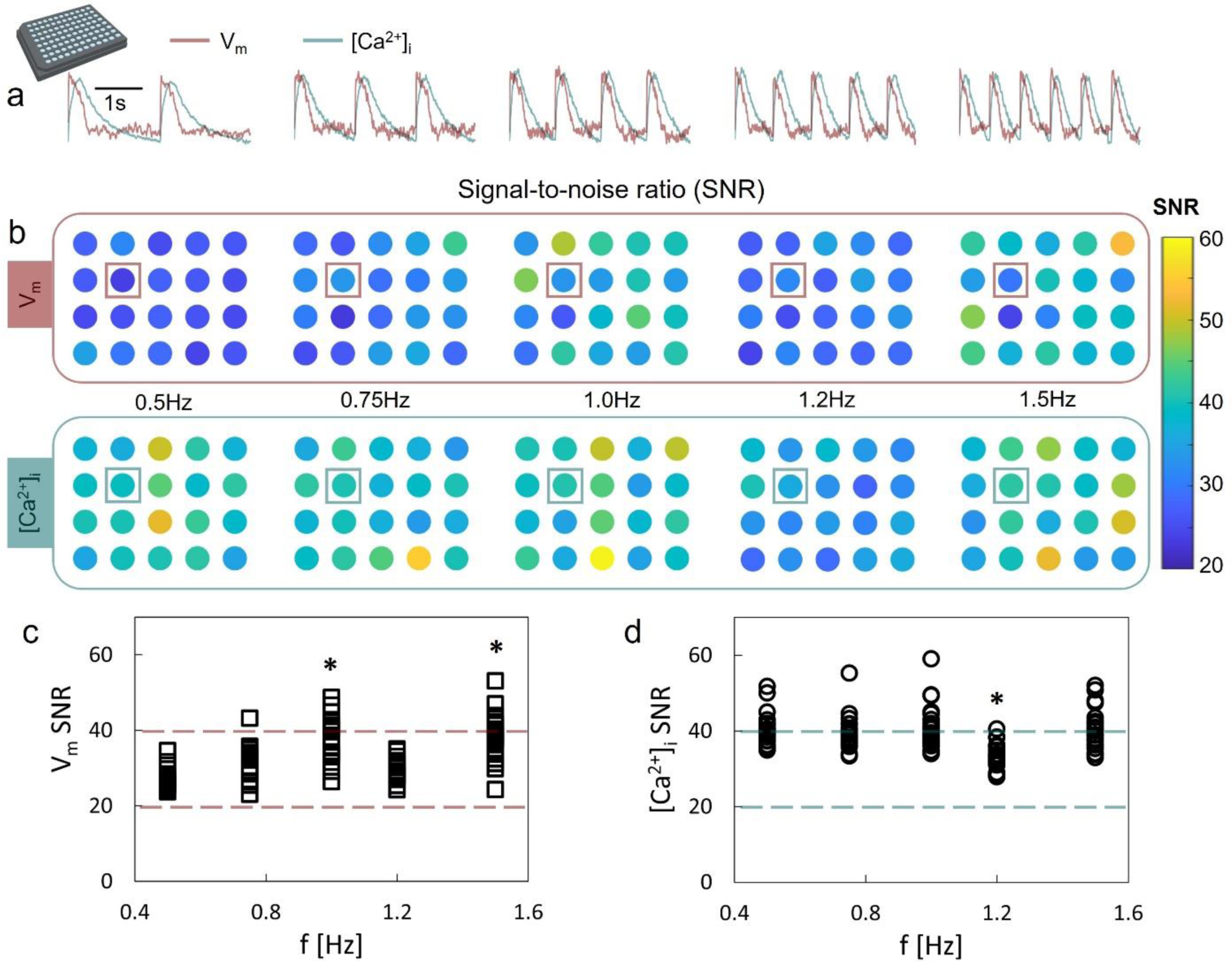
Signal-to-noise ratio (SNR) testing. HT plate imager testing using the central 20 wells of a 96 well plate. Optogenetically-modified syncytia of human induced pluripotent stem cell derived cardiomyocytes (hiPSC-CMs) were used. (a) Example V_m_ and [Ca^2+^]_i_ fluorescence traces for 0.5, 0.75, 1.0, 1.2, and 1.5Hz optical pacing; scale bar is 1s. The traces were from the same well, indicated with squares in (b). (b) Color coded SNR at each pacing frequency for both V_m_ (top panel) and [Ca^2+^]_i_ (bottom panel). (c) SNR of V_m_ records and (d) SNR of [Ca^2+^]_i_ as a function of of pacing frequency; n=20 biologically independent samples, two-way ANOVA. From the dashed lines, average SNR is approximately 30 for the V_m_ signals and 40 for the [Ca^2+^]_i_ signals.

Because pacing capture was successful in all 20 wells simultaneously, we concluded that the expanded beam of the single blue light source was sufficient for optogenetic pacing of the plate, allowing us to study restitution properties. **Figure 3a** shows an image of the 20 samples. From the V_m_ and [Ca^2+^]_i_ traces of the samples, we extracted various parameters: action potential duration at 80% repolarization, APD80, and calcium transient duration at 80% recovery, CTD80, as shown in **Figure 3b and c**, respectively, with their corresponding curve fittings. **Figure 3d** shows color-coded APD80 and CTD80 for each well at each pacing frequency. Variability among wells is relatively low, considering manual cell plating and labeling. Based on the good SNR and APD/CTD restitution characterizations, we therefore verified that the HT plate imager was fully functional for a large FOV – at least 37 x 45 mm (for the 20 sample) on a 96 well plate.

**Fig. 3.**
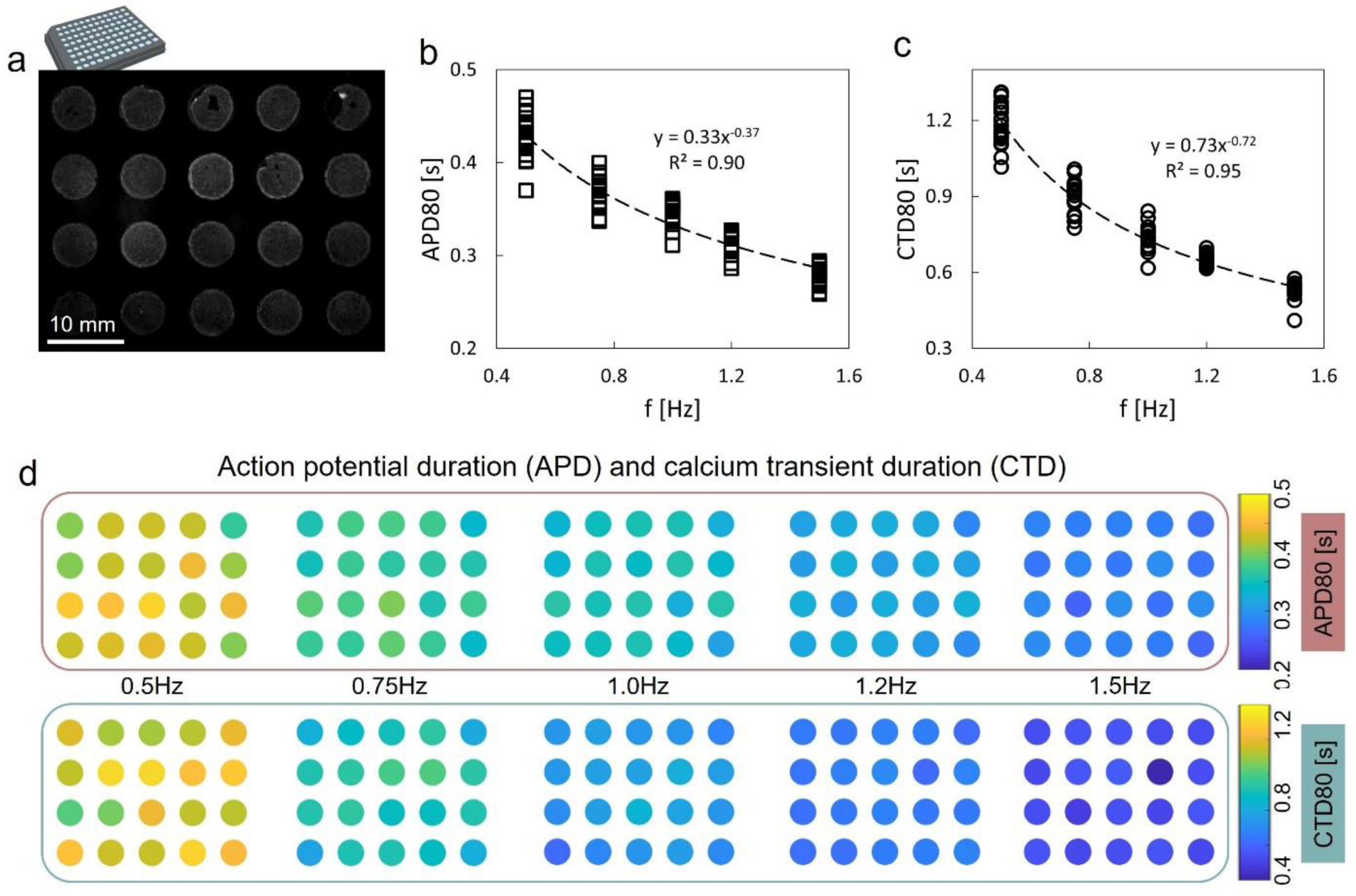
Parameter characterization and variability for voltage and calcium measurements. (a) An image of 20 hiPSC-CMs samples on a 96 well plate. Restitution plots of (b) action potential duration at 80% repolarization, APD80, and (c) calcium transient duration at 80% recovery, CTD80, with their corresponding curve fittings. (d) Color coded APD80 and CTD80 for the 20 wells at each pacing frequency; indicated color scales are in seconds.

### Drug testing using OptoDyCE-plate

The system was deployed to test the action of four drugs: dofetilide, nifedipine, cisapride and vanoxerine, compared to DMSO control samples. The drugs were applied acutely at concentrations, chosen to be 1 to 10x of the clinically used effective free therapeutic plasma concentrations (ETPC), **Table 1**. These drugs have varying degrees of pro-arrhythmia risk, as per Redfern’s classification[25]. Dofetilide (a class III antiarrhythmic) is a K+ ion channel blocker and identified as a high-risk drug. Nifedipine (anti-hypertension drug) is a blocker of the L-type calcium channel and a low-risk or safe drug. Cisapride is used as gastrokinetic agent and is classified as a medium risk drug. Vanoxerine[26–28] has been used for the treatment of cocaine addiction, as a multi-channel blocker, it was considered recently as an anti-arrhythmic agent[29–32], which unfortunately failed in phase III clinical trials due to cardiotoxic effects[33, 34].

**Table 1.** Drugs tested.

| Drug | Main use | Target(s) | ETPC | Concentration used | ETPC fold |
| --- | --- | --- | --- | --- | --- |
| **Dofetilide** | class III anti-arrhythmic | K<sup>+</sup> ion channel blocker | 2nM | 2nM | 1X |
| **Nifedipine** | anti-hypertensive | Ca<sup>2+</sup> ion channel blocker | 7.7nM | 77nM | 10X |
| **Cisapride** | gastrokinetic agent | serotonin 5-HT<sub>4</sub> receptor agonist | 2.6nM | 26nM | 10X |
| **Vanoxerine** | treatment of cocaine addiction | dopamine reuptake inhibitor | 100-700nM | 30, 100, 300, 1000nM | 0.3 to 10X |
| **TSA** | anti-fungal antibiotic | HDAC inhibitor (class I and II) | 100nM | 100nM | 1X |
| **Panobinostat** | chemotherapy (multiple myeloma) | pan-HDAC inhibitor | 10nM | 50nM | 5X |
| **SAHA (Vorinostat)** | chemotherapy (cutaneous T-cell lymphoma) | HDAC inhibitor (class I and II) | 0.3uM | 1uM | 3X |
| **4PB** | urea cycle disorders | HDAC inhibitor (class IIa) | 1mM | 5mM | 5X |
| **Tubastatin A** | anti-cancer | HDAC inhibitor (class IIb, HDAC6) | 1uM | 5uM | 5X |

Figure 4a shows functional responses (APD80 and CTD80) to the applied drugs in human iPSC-CMs during spontaneous or optically paced activity from five individual glass-bottom 96 well plates; Figures 4b-c illustrate the corresponding percentage change compared to control. We observed expected electrophysiological changes to the drug treatments, where prolongation of the APD is used as a measure of cardiotoxicity risk. At the applied concentrations, dofetilide, cisapride and vanoxerine induced frequency-dependent APD80 prolongation, while nifedipine-treated samples shortened APD80, as expected. For all four drugs, the effects were most pronounced at slower pacing rates, including during spontaneous activity, and the effects were significantly reduced at higher pacing rates, see **Table 2**. Two-way ANOVA showed significant effects of treatment and frequency on APD80 and CTD80, and a significant drug x frequency interaction. Individual drug responses were compared to control using post-hoc correction and the significant differences are indicated in Figure 4b-c. While calcium followed similar pattern of responses to that seen in voltage, the relative CTD80 changes under paced conditions were much smaller for these ion channel blockers and the CTD80 prolongation disappeared above 0.5Hz pacing. Despite plate-to-plate variability, the iPSC-CM responses to drugs were reproducible across plates. Dofetilide was used at low dose as a positive control, and even at this dose, some samples exhibited arrhythmic activity without pacing, as shown in the example traces, Figure 4b.

**Fig. 4.**
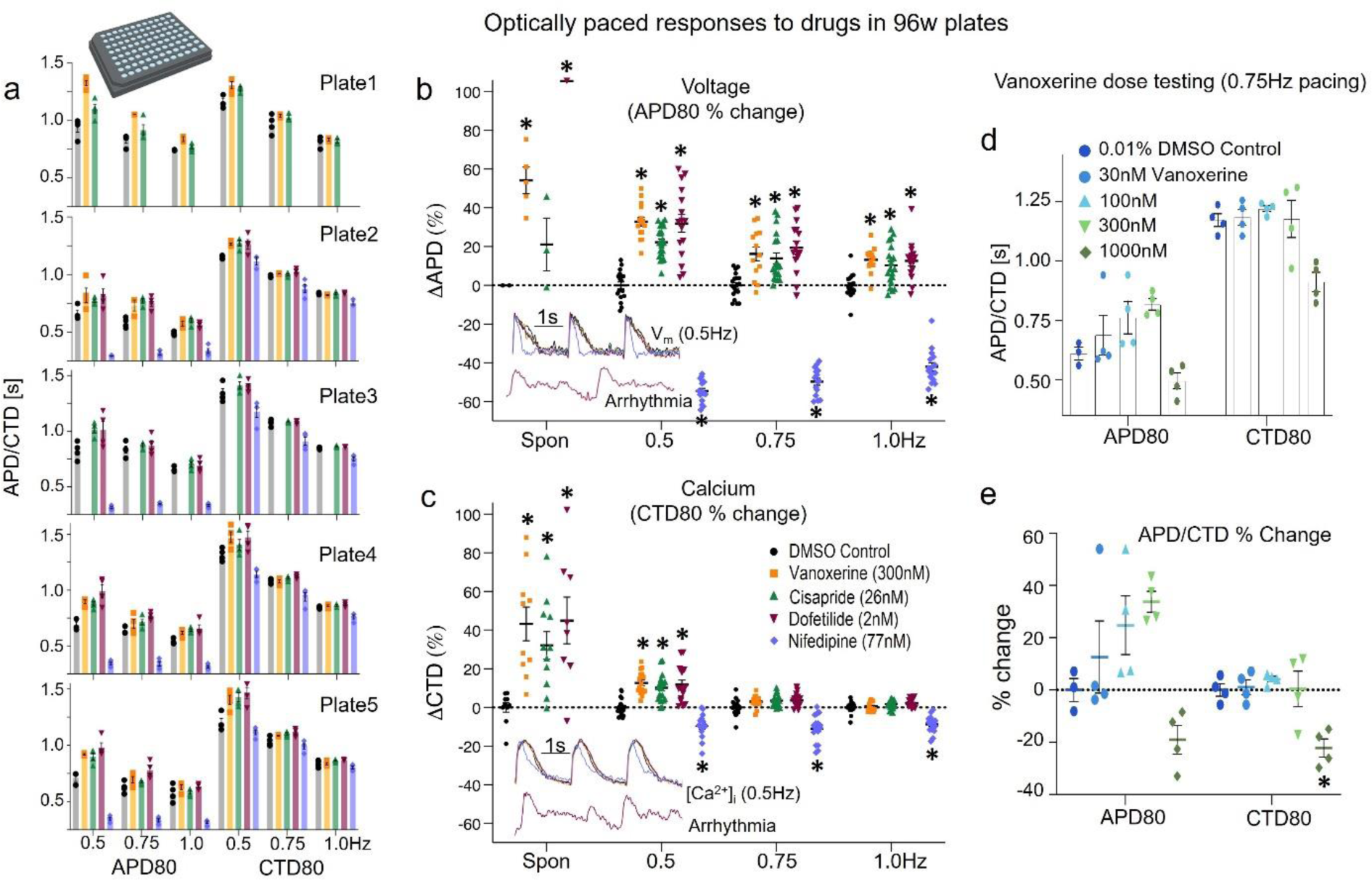
Drug testing in glass-bottom 96 well plates. Samples treated with vanoxerine (300nM), cisapride (26nM), dofetilide (2nM), and nifedipine (77nM) were compared with DMSO control (0.01 %) samples. (a) APD80 and CTD80 from five 96 well plates from different cultures under 0.5, 0.75, and 1.0Hz optical pacing. (b) Percentage changes in the corresponding APD80; nifedipine had no spontaneous activity and (c) percentage changes in CTD80 at each pacing frequency along with the values from the spontaneous activities (n=8-20 biologically independent samples per group; two-way ANOVA). Example V_m_ and [Ca^2+^]_i_ traces from all conditions at 0.5Hz optical pacing are included as insets, along with the arrhythmia traces from a dofetilide treated sample when no optical pacing was applied. The scale bar is 1s. (d) Dose response for vanoxerine (30, 100, 300, and 1000nM) compared with 0.01% DMSO control samples at 0.75Hz pacing (n = 3-4 biologically independent samples per group; one-way ANOVA). APD80 and CTD80 responses are shown. (e) Corresponding percentage changes in APD80 and CTD80. All error bars represent standard errors. In panels b and c, (*) indicate significance (at p<0.05) for the respective drug responses compared to control.

**Table 2.** Two-way ANOVA analysis on SNR as a function of pacing frequency and Vm vs. [Ca2+]i (corresponding to Fig. 2c, d).

| | pacing frequency | $V_m$ vs. $[Ca^{2+}]_i$ | freq x $V_m$ vs. $[Ca^{2+}]_i$ |
| --- | --- | --- | --- |
| SNR | **<0.0001** | **<0.0001** | **<0.0001** |

For vanoxerine, we tested a range of clinically relevant concentrations [28, 33, 34], Figures 4c-d. Expected dose-response effect in APD80 prolongation was seen for doses from 30 to 300nM, with little change in CTD80. At the highest dose of 1000nM, both APD80 and CTD80 collapsed to values below control. This can be explained as competition between vanoxerine’s effects on inward and outward currents, which have different kd for this drug. Overall, depending on dosing, vanoxerine can have quite variable effect on APD and some effect on CTD at high concentrations, **Table 3**. Monitoring calcium may not be a good indicator for cardiotoxic effects of vanoxerine.

**Table 3.** Two-way ANOVA analysis of drug responses under different pacing conditions in glass-bottom 96-well plates (corresponding to Fig. 4b, c).

|  | pacing frequency | drugs | freq x drugs |
| --- | --- | --- | --- |
| APD % change | **<0.0001** | **<0.0001** | **<0.0001** |
| CTD % change | **<0.0001** | **<0.0001** | **<0.0001** |

### HT interrogation of isotropic and anisotropic quasi-3D cell constructs

The conditions of syncytial cell growth may influence the functionality of human iPSC-CMs and their response to drugs. It is important to understand variations in the readouts due to microstructure differences in the cell constructs. Preserving the HT format (still using 96-well and 384-well plates), we explored quasi-3D cardiomyocyte assemblies on isotropic and anisotropic electrospun surfaces, Figures 5-6, with OptoDyCE-plate. The electrospun matrices within each well and the respective cardiomyocyte growth are shown in Figures 5a-b. The frequency-dependent functional responses from the HT plate imager on these surfaces are shown in Figures 5c-d, and as a comparison, testing on our microscope-based system from these samples is provided in Figure 5e. As seen both in the HT imager results, as well as in the microscope-based records, compared to the isotropic constructs, anisotropic cell growth yields hypertrophy-like responses, i.e., overall longer APDs and CTDs at baseline, and steeper restitution, corroborating previous results with primary rat cardiomyocyte syncytia[35, 36]. The effects of dofetilide, nifedipine and vanoxerine were tested, just like in the glass-bottom 96-well plates. Similar to the glass-bottom surfaces, these drugs exhibited stronger effects on voltage (compared to calcium) and these effects were more pronounced at lower pacing frequencies. The drug responses had tighter spread on the anisotropic surfaces for the same doses applied.

**Fig. 5.**
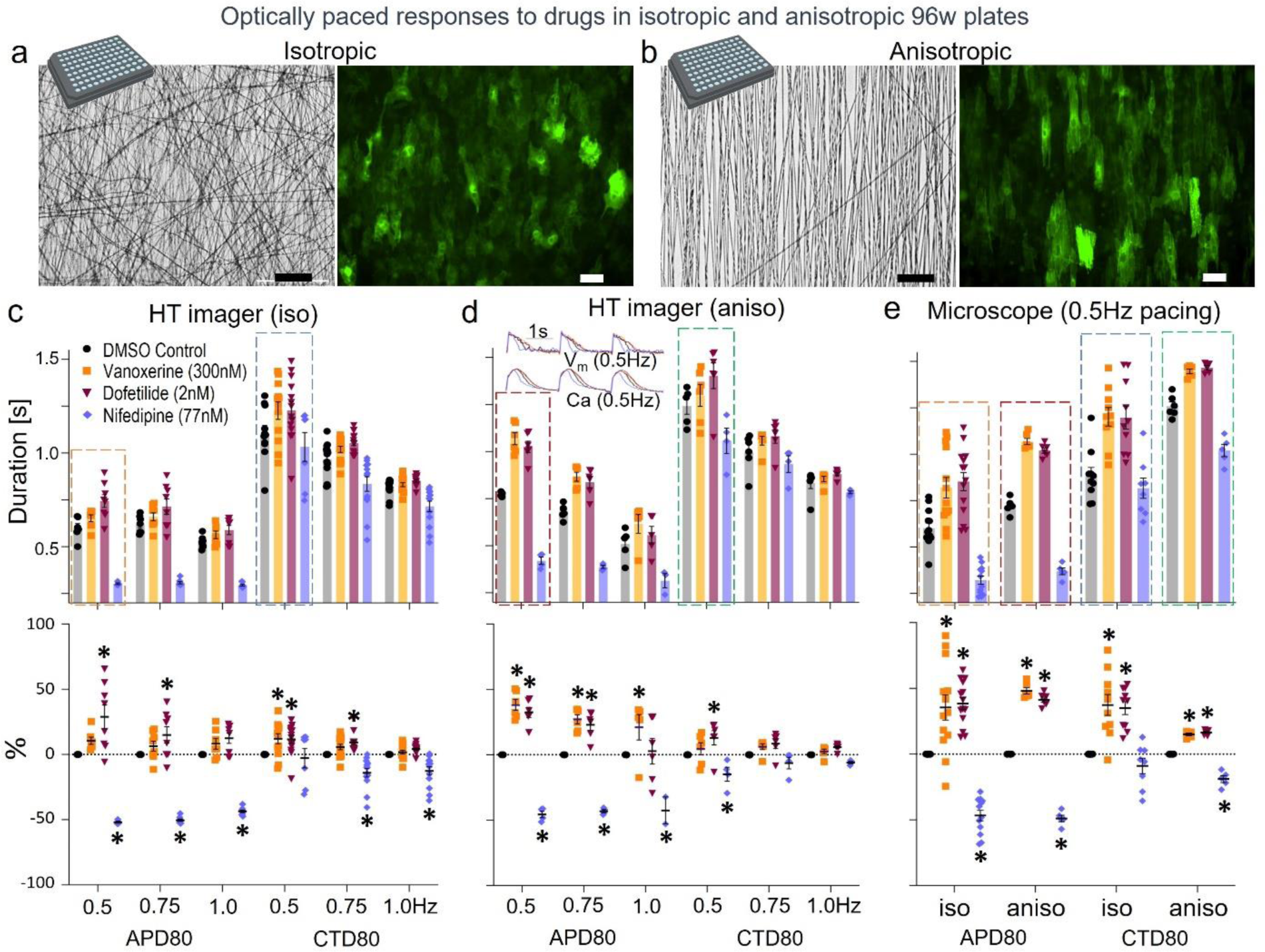
Drug testing in structured quasi 3D isotropic and anisotropic 96 well plates. (a) Image of the electrospun PCL for an isotropic well in 96 well plate and the corresponding image of cell growth in isotropic sample (green is ChR2-eYFP). (b) Similar images for an anisotropic electrospun sample in 96 well format. Scale bar is 50μm. (c) APD80 and CTD80 responses to drugs from the HT full plate imager in isotropic samples in 96 well plate under optical pacing with corresponding percentage changes, (n = 4-14 biologically independent samples per group; two-way ANOVA). (d) Same as (c) for anisotropic samples with corresponding percentage changes, (n = 2-6 biologically independent samples; two-way ANOVA). Insets show example voltage and calcium traces. (e) Microscope-based APD80 and CTD80 measurements for 0.5Hz pacing in isotropic and anisotropic samples with corresponding percentage changes, (n = 5-14 biologically independent samples per group). The error bars represent standard errors. In panels c-e, (*) indicate significance (at p<0.05) for the respective drug responses compared to control.

**Fig. 6.**
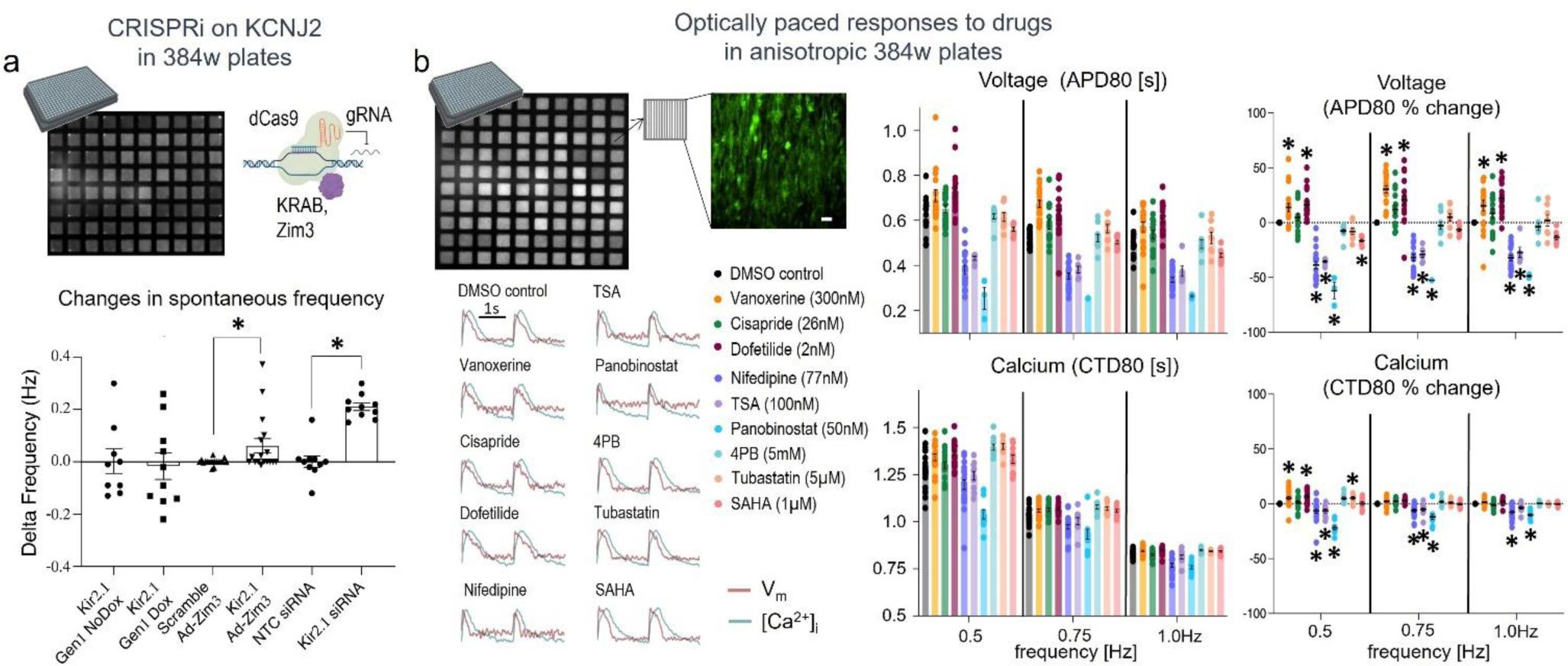
HT plate imager deployed for 384 well plate measurements. (a) CRISPRi and siRNA testing in glass-bottom 384 well plates, using different effectors and parallel measurement of spontaneous activity from 80 samples. The bottom panel presents a comparison of the relative changes in spontaneous frequency from control using different genetic perturbation methods targeting KCNJ2/Kir2.1 (n = 9-18 biologically independent samples per group; unpaired t-test). (b) Drug testing in quasi-3D anisotropic samples in 384 well plate – parallel measurements of voltage and calcium from 100 samples. Shown is an image of the 100 samples, an image of the aligned cells (scale bar: 50 μm) in one well, and example V_m_ and [Ca^2+^]_i_ traces from each of the conditions. The right panel shows APD80 and CTD80 changes upon drug treatments of the hiPSC-CMs under different optical pacing with corresponding percentage changes (n = 8-10 biologically independent samples per group; two-way ANOVA). The error bars represent standard errors. Significance (at p<0.05) for the respective responses to treatments compared to control is indicated by a (*).

Two-way ANOVA analysis indicated that drug treatment and pacing both caused significant changes in the anisotropic and in the isotropic samples in voltage and calcium, when tested with the HT plate imager and when tested on the microscope-based system, **Tables 4-6**. The interaction term (drug x frequency) was only significant in the microscope-based data for action potential duration. Despite similar trends to the APD80/CTD80 prolongation and shortening to the results from Figure 4, the pairwise comparisons to control, after posthoc correction, in the isotropic samples did not reveal differences for vanoxerine but yielded significant APD80 changes for dofetilide and nifedipine, when measured with the plate imager, Figure 5c. In the anisotropic samples, vanoxerine had a significant effect on APD80 across all pacing frequencies; dofetilide also prolonged APD80 at 0.5 and 0.75 Hz, and CTD80 at 0.5 Hz pacing. Nifedipine had a significant effect on APD80 for each pacing frequency and on CTD80 at 0.5Hz pacing. For all tested configurations (monolayers on glass, quasi-3D samples grown isotropically and anisotropically), the drug effects were qualitatively similar, with expected trends in APD and CTD, Figures 4 and **5**. In Figure 5, we have validated the results from the full-plate imager with sequential microscopic OptoDyCE imaging at 0.5Hz optical pacing. The durations obtained from the imager and the microscope agree with each other.

**Table 4.** Two-way ANOVA analysis of drug responses in isotropic quasi-3D electrospun 96-well plates (corresponding to Fig. 5c).

|  | pacing frequency | drugs | freq x drugs |
| --- | --- | --- | --- |
| APD | **<0.0001** | **<0.0001** | 0.074 |
| CTD | **<0.0001** | **<0.0001** | 0.5291 |

**Table 5.**
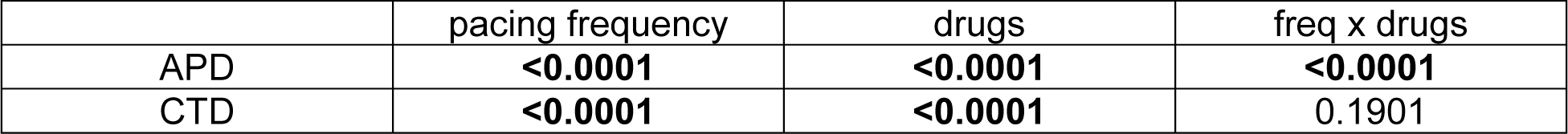
Two-way ANOVA analysis of drug responses in anisotropic quasi-3D electrospun 96-well plates (corresponding to Fig. 5d).

**Table 6.** One-way ANOVA analysis of drug responses in isotropic and anisotropic plates 96-well plates using microscope-based measurements at 0.5Hz pacing frequency (corresponding to Fig. 5e).

|  | Van vs. control | Dof vs. control | Nif vs. control |
| --- | --- | --- | --- |
| Iso APD | **0.0018** | **0.0002** | **<0.0001** |
| Iso CTD | **0.0004** | **0.0004** | >0.9999 |
| Aniso APD | **<0.0001** | **<0.0001** | **<0.0001** |
| Aniso CTD | **<0.0001** | **<0.0001** | **<0.0001** |

### Validation of HT testing in 384-well plates

OptoDyCE-plate was also deployed for multimodal parameter measurements on 384 well plate samples. Figure 6a illustrates measurements from 80 samples (8 x 10) in glass-bottom 384-well plate. Utilizing interference CRISPR (CRISPRi) and siRNA, targeting KCNJ2 in the hiPSC-CMs, we tested 8 conditions with 10 samples for each. Two different effectors for CRISPRi were tested (Dox-inducible system with dCas9-KRAB[37] and a constitutive expression system with dCas9-Zim3[38]), as described in [39]. In addition to the conditions listed in Figure 6a, there was a group without any perturbation and a group with dCas9-Zim3 without gRNA. As KCNJ2 encodes for the inward rectifier current, IK1, responsible for the stability of the resting membrane potential, our output parameter, was spontaneous beating frequency, based on calcium records quantified using OptoDyCE-plate. From the results, the inducible dCas9-KRAB targeting KCNJ2 in high density syncytia was not effective in altering spontaneous beat frequency, as we have shown in[39]. The improved CRISPRi-based system (dCas9-KRAB-Zim3) did have a more pronounced effect on spontaneous beat frequency when targeting KCNJ2. The changes in spontaneous beat frequency were most reliable when using siRNA, **Table 7**. Overall, the 384-well format allows for quick testing of multiple conditions when developing new genetic modification tools.

**Table 7.** Unpaired t-test between the treatments to suppress KCNJ2: Dox/noDox, Zim3+/−, and siRNA +/−, for spontaneous activities of iPSC-CMs in glass-bottom 384-well plates (corresponding to Fig. 6a bottom panel).

| Kir2.1 Gen1 Dox vs. noDox | Scramble vs. Kir2.1 Ad-Zim3 | NTC vs. Kir2.1 siRNA |
| --- | --- | --- |
| 0.816 | 0.0835 | **<0.00051** |

In Figure 6b, we applied OptoDyCE-plate in 100 samples (10 x 10) in 384-well plate with anisotropic quasi-3D cell growth to test an extended list of pharmacological agents, as described in **Table 1**, including five histone deacetylase inhibitors (HDACi). These HDACi were panobinostat, trichostatin-A (TSA), 4-phenylbutyrate (4PB), tubastatin and SAHA (Vorinostat), applied chronically for 48 hours prior to measurements. HDACi have global effects on the gene expression of multiple genes and their effects on cardiac electrophysiology and cardiotoxicity are under-studied[40–44].

The SNR obtained in these 384-well samples was sufficient for quantification of parameters. The measured APD80, CTD80 and the drug effects were consistent with the ones obtained in anisotropic 96-well plates, as in Figure 5. The APD80 changes for these anisotropic samples in response to dofetilide, vanoxerine and nifedipine (Figure 6b) were more pronounced than the ones seen in glass-bottom dishes (Figure 4), although cisapride showed only slight APD80 prolongation that was not significant. Drug application and pacing independently and as a cross-term were found to be significant modulators of APD80/CTD80, **Table 8**. From the HDACi applied in these samples, the most pronounced effects were seen for the pan-HDAC inhibitor, panobinostat, which shortened both APD80 and CTD80, in a frequency-dependent manner. Clinically, adverse electrophysiological effects have been reported previously for panabinostat[41]. SAHA (vorinostat) also shortened the APD80 significantly at low pacing frequencies. TSA reliably shortened APD80 at all tested pacing rates and had milder effect on CTD80. Conversely, at the applied concentrations, 4PB and Tubastatin yielded small but significant (for Tubastatin) CTD80 prolongation at low pacing rates, despite no effect on APD80. These are examples of differential effects on voltage and calcium that some drugs can exert, and the need for comprehensive assessment of electromechanical function. The HDACi cardiotoxicity may not fit some of the classic measures based on HERG channel block, leading to APD prolongation and increased risk for Torsade-de-pointes (TdP) type of arrhythmias. In our experience and as shown here, in most cases, HDACi led primarily to APD80 shortening.

**Table 8.** Two-way ANOVA analysis of drug responses in anisotropic quasi-3D electrospun 384-well plates (corresponding to Fig. 6b right panel).

|  | pacing frequency | drugs | freq x drugs |
| --- | --- | --- | --- |
| APD | **<0.0001** | **<0.0001** | 0.1019 |
| CTD | **<0.0001** | **<0.0001** | **<0.0001** |

Overall, using OptoDyCE-plate in the 384-well plate format allows the comprehensive testing of multiple samples and conditions simultaneously, under different pacing rates. The FOV for the 100 samples was 46 x 46 mm, and based on illumination profile, likely could be extended to optically stimulate and image up to approximately 200 samples in these plates.

### Optical point pacing through ChR2 spheroids integrated in 96-well samples

The action of some drugs has unique effects on conduction properties rather than the functionality at the single-cell level. Therefore, it is desirable to be able to study wave propagation in a HT format. In preliminary investigation, we explored the possibility to create an optical point stimulation in each well, without patterned light, using the simple design of OptoDyCE-plate. To do so, we utilized our tandem-cell-unit approach[45] for optogenetic pacing using ChR2-HEK (spark) spheroids[46] deposited on the iPSC-CM monolayers, Figure 7. Activation maps and traces for both V_m_ and [Ca^2+^]_i_ from three samples (S1, S2, and S3) illustrate the responses at 0.5, 0.75, and 1.0Hz optical pacing. Note that the duration of optical pacing was 20ms to excite the spheroids. As expected, the waves for each sample originated from approximately the same location (of the spheroid) at each pacing frequency. From the activation maps, conduction velocities (CVs) can be obtained in a high-throughput manner. In these examples, we did not observe expected CV restitution (CV slowing with increased pacing frequency), as normally seen in larger paced syncytia. It is unclear if the small size and particular geometry of the 96-well samples may be the reason for this. Optimization and automated deposition of the spark-cell spheroids can enable future studies of conduction abnormalities in a HT format using OptoDyCE-plate, without optogenetically modifying the cardiomyocytes. Higher acquisition rate will further increase the range of measurable CVs in the small samples in 96-well format.

**Fig. 7.**
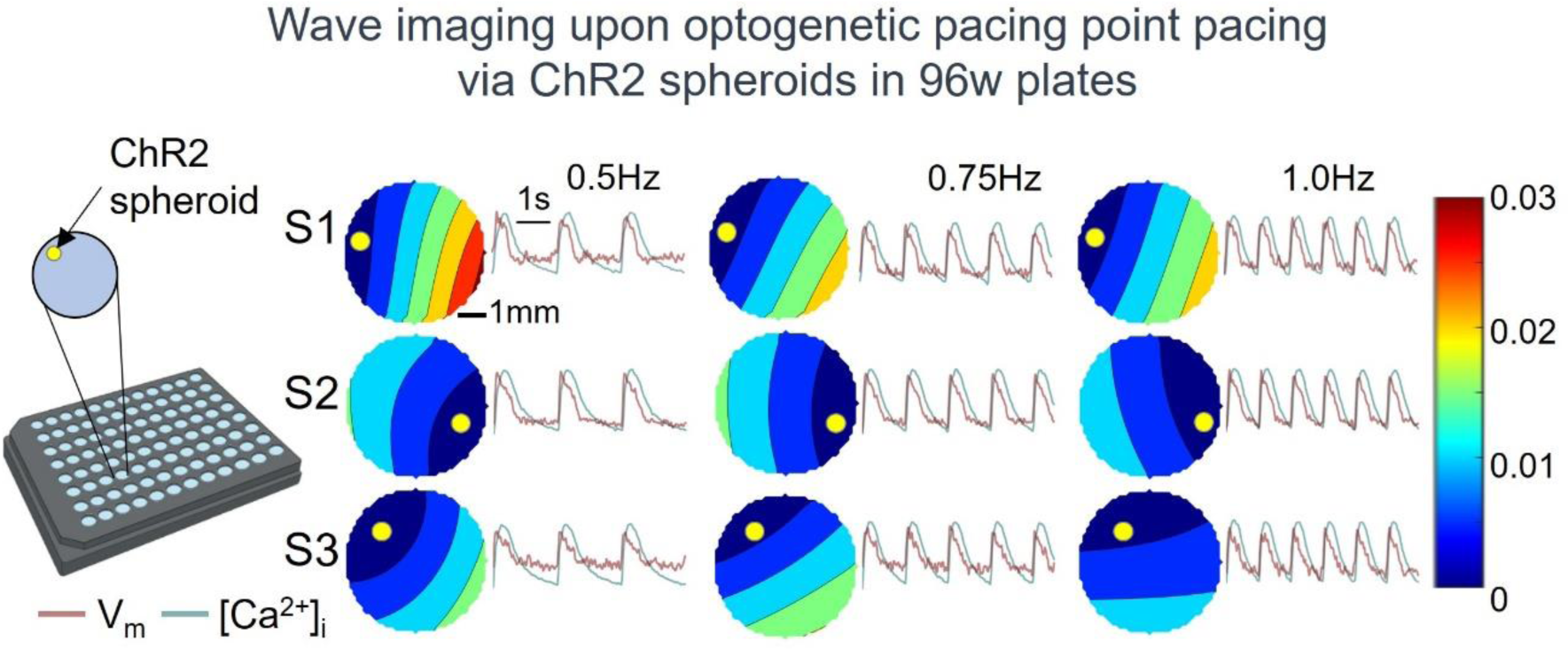
HT plate imager deployed for wave imaging with optogenetic pacing via integrated ChR2 spheroids on unmodified iPSC-CM monolayers in glass-bottom 96 well plates. The left panel illustrates a sample. Shown are activation maps and traces for both V_m_ and [Ca^2+^]_i_ from three samples at 0.5, 0.75, and 1.0Hz pacing, with scale bars indicating 1s and 1mm and a color bar indicating the wavefront activation times (in seconds). Yellow markers indicate the positioning of the spheroid for each of the samples.

### Validation of temporal multiplexing for single-sensor multiparametric imaging

All the experiments above were conducted when either red or green LED was constantly on for V_m_ or [Ca^2+^]_i_ recording, respectively, in a sequential manner in the same samples. This way, V_m_ and [Ca^2+^]_i_ were recorded at two different times (up to a minute apart). To further advance the system, we then implemented a temporal multiplexing scheme where both V_m_ are [Ca^2+^]_i_ were recorded simultaneously using the same low-cost camera and fast LED switching, synchronized with the camera frames, as described in our previous microscope-based system[12]. Figure 8a shows the scheme of the temporal multiplexing. Here, we used Arduino 2560 to control both LEDs and the camera. During one frame, the red LED is on, and the green LED is off, for the next frame, the red LED is off, and the green LED is on. The camera operated at 100 frames per second (fps) and recorded V_m_ during one frame and [Ca^2+^]_i_ the next frame, and so on, while post-processing compiled the voltage and calcium recordings from the interlaced frames. Note that optical pacing events were not necessarily synchronized with the camera or the green and red LEDs. The duration of optical pacing was 10ms. We tested the temporal multiplexing mode on samples from a 96 well plate, with 0.5, 0.75, and 1.0Hz optical pacing frequencies. Figure 8b presents the restitution plots for APD80 and CTD80. Open symbols are the values obtained with temporal multiplexing scheme. For comparison, the APD/CTD durations were also obtained with either LED constantly on, shown as solid symbols. The two schemes agreed well with each other. Figure 8c-d show example traces from temporal multiplexing and constant illumination, respectively, at each pacing frequency. SNR was similar for constant illumination vs. multiplexing – approx. 30 to 40 for both V_m_ and [Ca^2+^]_i_.

**Fig. 8.**
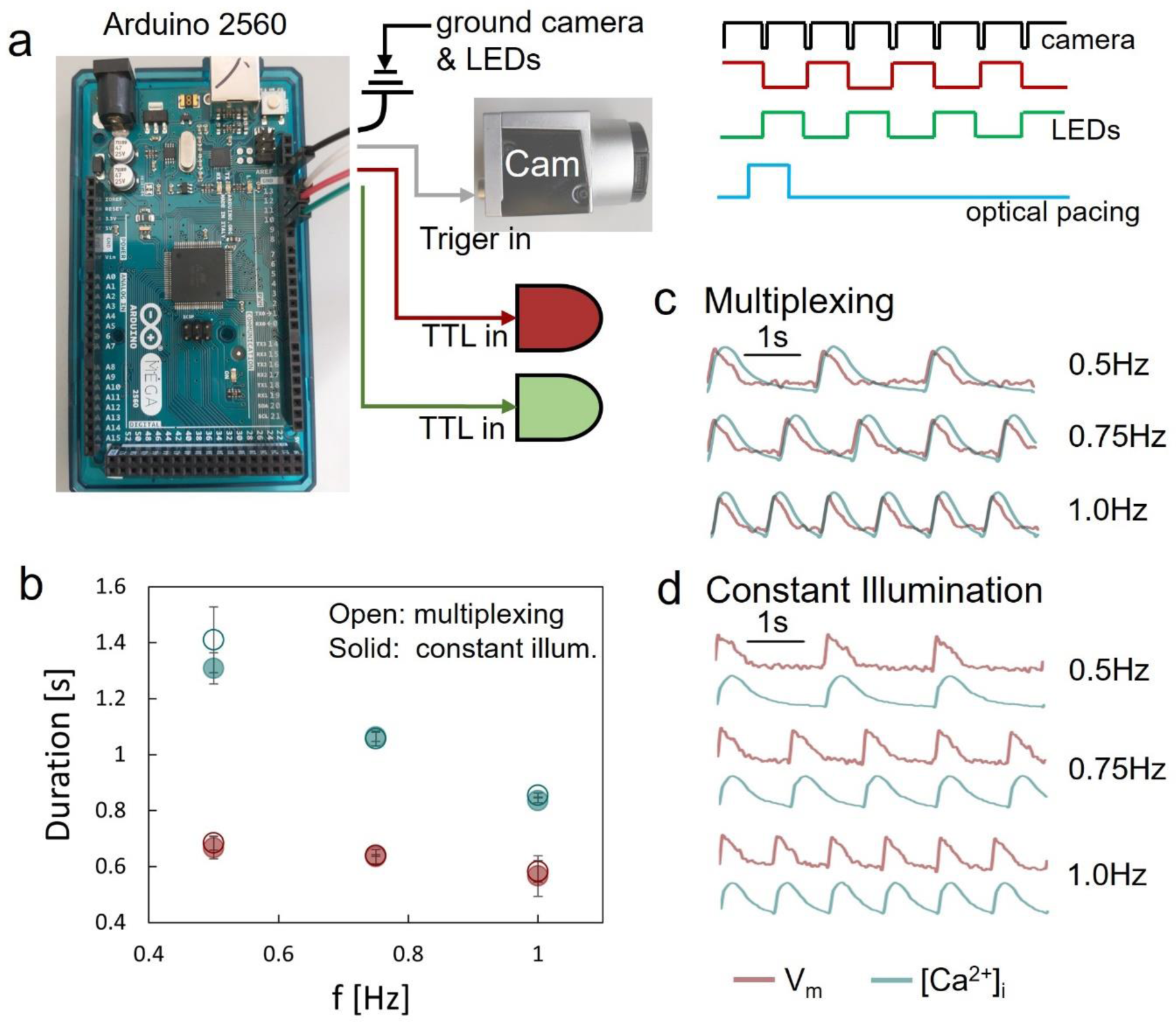
Validation of temporal multiplexing for simultaneous voltage and calcium measurements with the low cost camera of the HT plate imager. (a) Diagram of Arduino Mega 2560 controlling circuit. Green and red LEDs are multiplexed. Each frame of the camera records when only one LED is on and the other is off. Camera runs at 100 fps with exposure time 9.8ms for each frame. Optogenetic pacing uses 10ms pulses, not necessarily aligned with each time frame. (b) Restitution plots for APD80 and CTD80 (n = 4∼6). Open symbols are the values obtained from temporal multiplexing scheme and solid symbols are from constant illumination. Error bars represent standard deviation. (c) Example V_m_ and [Ca^2+^]_i_ traces obtained simultaneously from temporal multiplexing at each pacing frequency. (d) Example V_m_ and [Ca^2+^]_i_ traces obtained separately with constant illumination at each pacing. The scale bar indicates 1s duration.

## Methods

### Building Opto-DyCE96 full plate imager for all-optical electrophysiology

The full high throughput (HT) plate imager for all-optical electrophysiology is structured on a 12” x 12” aluminum breadboard; all mechanical components are standard parts from Thorlabs, Newton, NJ, except for the filter holder-adaptor unit (Edmund Optics, Barrington, NJ) attached to the main imaging lens. Based on epi-illumination for the multi-modal imaging of V_m_ and [Ca^2+^]_I_, fluorescence signals were simultaneously obtained in a full plate under optical/optogenetic stimulation. Figure 1a and b show the schematic diagram and photo of the system, which uses a low-cost camera, Basler acA720-520um (Basler AG, Germany), with 720×540 pixels, 6.9µm per pixel, capable of running at up to 525 frames per second (fps). The camera collects the emitted fluorescence to image V_m_ and [Ca^2+^]_i_. The imaging lens is a 12mm camera lens (f/2.0, Thorlabs, Newton, NJ). The combination of light sources and multi-band filters is previously described[12, 17]. In the current implementation, the light sources of the system are high-power LEDs: a red LED (SOLIS-660C, Thorlabs) at 660nm for excitation of the voltage-sensitive dye and a green LED (SOLIS-525C, Thorlabs) at 530nm for excitation of the calcium-sensitive dye. The red and green LEDs are paired with excitation filters: F1 and F2 (AT655/30m and ET525/30m, Chroma, Bellows Falls, VT), respectively, and combined by a 605 long pass dichroic filter F3 (FF605-DI02-50.8X72.0, Semrock, Rochester, NY). Optogenetic stimulation is achieved via a high-power blue LED (SOLIS-470C, Thorlabs) at 470nm, typical pulse duration 10ms, controlled by TTL signals via an LED driver. The blue LED is equipped with an excitation filter F4 (ET470/24m, Chroma), and combined by a 495 long pass dichroic filter F5 (FF495-DI03-50.8X72.0, Semrock) to join the red and green lights. The three lights are then reflected by a multiband dichroic filter F6 (ZT473/532/660rpc-xt-UF1, Chroma) and propagate towards the sample from below. The emitted V_m_ and [Ca^2+^]_i_ fluorescence signals are collected by the same multiband dichroic filter F6, the multi-bandpass emission filter F7 (ET595/40m + 700lp, Chroma), and the camera lens attached to the camera. Since the high-power LEDs are large, instead of the standard sizes of the filters used in [17], all filters applied in the HT plate imager are enlarged accordingly. All dichroic filters are mounted in kinematic cubes (DFM2, Thorlabs). Basler Video Recording Software was used for constant illumination recordings and Pylon Viewer for temporal multiplexing illumination recordings using a single camera.

### HT 96-well and 384-well plates

For HT studies, we used 96-well glass-bottom plates (Cat. P96-1-N, Cellvis) and 384-well glass bottom plates (Cat. P384-1.5H-N, Cellvis). For studies of structured quasi-3D cell growth, plates with anisotropic and isotropic electrospun fiber arrangement (700nm polycaprolactone or PCL fibers) were obtained from Nanofiber Solutions (SKU 9601, 9602, 38401, 38402).

### hiPSC-CM cell preparation

iCell Cardiomyocytes^2^ (Cat. C1016, Donor 01434, Fujifilm/Cellular Dynamics) were cultured onto fibronectin (Cat. 356009, Corning) coated (50µg/mL) HT 96-well or 384-well plates at 1.56 x 10^5^ hiPSC-CMs cells/cm^2^ and maintained according to the manufacturer’s protocol. For experiments conducted on 384-well electrospun fiber plates, cells were plated at 2.75 x 10^5^ cells/cm^2^. All hiPSC-CMs were mycoplasma tested and cultured at 37°C and 5% CO_2_.

### Spark-cell spheroids and iPSC-CM monolayer integration

As per Chua et al[46], WT HEK cells were cultured using Dulbecco’s Modified Eagle Medium (DMEM) with 10% fetal bovine serum (FBS) and 1% penicillin-streptomycin. To express optical actuator ChR2, WT HEK cells were transduced with Ad-CMV-hChR2(H134R)-eYFP (Vector Biolabs) and magnetic nano-particles (Cat. 657841, Greiner BIO-ONE) were added and cultured overnight for complete absorption. The following day, the cells were trypsinized and 30,000 cells/spheroid were plated onto Corning 96-well spheroid microplates (Cat. CLC4920, Millipore Sigma) for 24 hours for spheroid aggregation. The ChR2-expressing spheroids were pulled down onto the hiPSC-CM monolayer by a magnetic plate and integrated for up to 24 hours prior to functional experiments.

### CRISPRi gene modulation

For targeting the potassium channel gene KCNJ2 in the HT knockdown studies, we used CRISPRi as previously described[37, 39]. Guide RNA’s were designed using CRISPR-ERA (Stanford) and VectorBuilder’s sgRNA design software to target around the transcription start site (TSS). gRNA oligo sequence ATTCCCAAGACCCAGCCCGC targeting KCNJ2 and gRNA oligo GTGTAGTTCGACCATTCGTG for Scramble were cloned into a lentiviral vector and packaged into lentiviruses by VectorBuilder.

For CRISPRi knockdown of KCNJ2 by Doxycycline (Dox)-inducible CRISPRi (with KRAB effector domain), cells were transduced with both gRNA lentivirus and then transfected hiPSC-CMs with 0.022µg AAVS1-Talen-L, a gift from Danwei Huangfu (Cat. 59025, Addgene), 0.022µg AAVS1-Talen-R, a gift from Danwei Huangfu (Cat. 59026, Addgene) and 0.044µg pAAVS1-Ndi-CRISPRi, a gift from Bruce Conklin (Cat. 73497, Addgene) per 17 x 10^3^ hiPSC-CMs (Mandegar et al. 2016) using Lipofectamine 3000 transfection agent (Cat. L3000001, ThermoFisher) following the manufacturer’s protocol. 2 µM of Dox (Cat. D9891, Sigma-Aldrich) was applied for 5 days for full expression.

For CRISPRi knockdown of KCNJ2 by dCas9-KRAB-Zim3[38], we packaged pHR-UCOE-SFFV-dCas9-mCherry-ZIM3-KRAB, a gift from Mikko Taipale (Cat. 154473, Addgene) into adenovirus Ad-dCas9-mCherry-Zim3 through VectorBuilder. Human iPSC-CMs were transduced with both gRNA lentivirus and Ad-dCas9-mCherry-Zim3 at MOI 500 and co-labelled with voltage- and calcium-sensitive dyes for functional experiments 5 days post transduction.

### siRNA gene modulation

For knockdown of KCNJ2 by siRNA, cells were transfected with 270ng of esiRNA Human KCNJ2 (Cat. EHU019661, Millipore Sigma) or non-targeting control (NTC) siRNA (Cat. SIC001, Millipore Sigma) per 2.7 x 10^5^ hiPSC-CMs 5 days post plating using Mission Transfection Reagent (Cat. S1452, Millipore Sigma) following the manufacturer’s instructions[39, 47]. Cells were labelled for functional experiments 4 days post-transfection.

### Drug treatment and testing

All drugs were obtained from Sigma Aldrich: vanoxerine (Cat. D052), cisapride (C4740), dofetilide (PZ0016), nifedipine (N7634), as well as the following histone deacetylase inhibitors, HDACi: 4-phenylbutyrate, 4PB (567616), tubastatin A (SML0044) and from a kit (EPI009-1KT): trichostatin-A (TSA), panobinostat, and SAHA. The drugs were dissolved in DMSO (Cat. 34869, Sigma-Aldrich), except for 4PB (dissolved in sterile water) and stored in −20°C until ready for application on the hiPSC-CMs. The HDAC inhibitors TSA, panobinostat, 4PB, tubastatin, and SAHA stock solutions were diluted in hiPSC-CM maintenance media and applied to the cells 48 hours prior to functional measurements. Vanoxerine, cisapride, dofetilide, and nifedipine stocks were diluted in Tyrode’s solution on experimental day and applied for 30 minutes at 37°C, 5% CO2 prior to functional experiments. Drug concentrations are as specified in the figures, and all were selected to be at 1X to 10X of the effective free therapeutic plasma concentrations, ETPC, see **Table 1**.*All-optical functional experiments*.

For all-optical electrophysiological studies, hiPSC-CMs were infected with Ad-CMV-hChR2(H134R)-eYFP (Vector Biolabs) at least two days prior to experiments for optical actuation. Membrane voltage and intracellular calcium staining protocol was performed as previously described [12, 13]. All experiments were conducted at 32-35°C in Tyrode’s solution (mM): NaCl, 135; MgCl2, 1; KCl, 5.4; CaCl2, 1.8; NaH2PO4, 0.33; glucose, 5; and HEPES, 5 at pH 7.4. Optical recordings of voltage were performed using the synthetic dye BeRST1 diluted in Tyrode’s solution at 1 µM and calcium was recorded using Rhod-4 diluted in Tyrode’s solution at 10 µM. All probes are spectrally compatible with ChR2.

### Statistical analysis

Biological replicates are described in the figure legend. Statistical analysis was conducted in Prism Software (GraphPad). One-way ANOVA followed by post-hoc Bonferroni tests were used to evaluate significance where a gradient of vanoxerine dosages were used on the hiPSC-CMs. Two-way ANOVA followed by post-hoc Bonferrroni tests were used to evaluate significance between samples with varying pacing frequencies and drug treatments, as well as when comparing SNR of V_m_ and [Ca^2+^]I at varying frequencies. Finally, an unpaired t-test was used to assess the significance between spontaneous beat rate of hiPSC-CMs that have been genetically modified to target KCNJ2. Significant differences were reported for p < 0.05.

## Discussion

Opto-DyCE-plate provides a simple, affordable, and versatile solution to high-throughput drug screening for cardiotoxicity testing and drug development. The total cost of our system is less than $20,000, lower than the price of a single scientific camera. To our knowledge, this is the first low-cost highly parallel system of this kind to offer multiparametric electromechanical assessment in >100 samples simultaneously over a large FOV, in standard HT plates (96- and 384-well) used in the pharmaceutical industry. While HT solutions based on planar patch clamp[48], including such covering full-plate format, have evolved rapidly over the last decade, their inherent limitation lies in the need for contact and the use of isolated cells, which may not capture truthfully the cellular properties and responses within a cardiac syncytium[49, 50]. Another HT approach, the microelectrode arrays, MEAs, has also seen rapid development and has been adopted to actuation and interrogation of cellular monolayers grown in more expensive specialized HT plates[51, 52]. This approach is attractive as a long-term tool due to its label-free nature, however extracellular potentials captured by MEAs are acutely dependent on good contact and their interpretation for variable action potential shapes is non-trivial, i.e. they may not capture well key features like APD80. Overall, compared to other competing approaches, all-optical electrophysiology[12, 13, 16, 17, 53] offers uniquely comprehensive and robust methods for stimulation and multiparametric recordings within syncytial structures without direct contact. As we show here, the implementation can be very low cost and the consumables can be standard HT plates.

In validating the deployment of OptoDyCE-plate with human iPSC-CMs, we tackled several important questions:

- Does pacing rate matter in the assessment of responses to drugs?
- Are voltage and calcium measurements always synonymous in assessing cardiotoxicity, and can calcium transients be used as a surrogate measure for electrical abnormalities?
- Does the cell growth setting (monolayers vs. quasi-3D isotropic or anisotropic growth) alter the predicted responses to drugs?

The answers to these questions inform us how future efforts should be directed to provide simple yet reliable preclinical drug testing.

In the presented OptoDyCE-plate system, optogenetic pacing (global and point-pacing) adds an important feature compared to other systems for optical recordings of spontaneous activity[7, 20, 54], namely control over rate to better quantify drug responses. In all our studies, pacing frequency was not only an independent modulator of voltage and calcium morphology, but also an important factor in the cardiac response to drugs. Reverse use dependence (stronger response to drugs at slower rates)[55] is common for drugs that affect the HERG channel and therefore prolong APD. This stronger drug response at spontaneous activity or lower pacing frequency was indeed seen for most of the compounds tested in this study, Figures 4-6. However, there are perturbations that are better revealed at higher pacing rates, such as drugs affecting conduction velocity, for example, as seen in other studies[39]. Rate control is essential in quantifying drug action accurately, as illustrated in the presented results here.

Fluorescence based measurements of intracellular calcium have been attractive for HT imaging because of the superior quality of optical sensors for calcium (compared to those for voltage), and some commercial systems are based on such measurements[7, 56]. These are often used as direct surrogate for quantitative cardiotoxicity studies. Conversely, MEA-based records exclusively rely on extracellular measurements to assess potential drug effects, while missing potentially important effects on calcium. Very few studies have provided detailed view of the similarities and differences in voltage and calcium responses to drugs. Based on our data, if calcium-based measurements were used to assess the effects of a drug like vanoxerine on cardiac electrophysiology, they would miss a significant APD80 prolongation at low to intermediate clinically relevant doses, Figure 4. At higher pacing rates, calcium-only assessments may also miss or underestimate the action of classic drugs known to modulate cardiac electrophysiology, even including dofetilide, Figures 4-6. Alternatively, some compounds may have a preferential effect on the calcium transients without detectable APD modulation, as for example the HDACi tubastatin at low pacing rates, Figure 6b. Overall, we argue that in many cases calcium may not be a suitable surrogate for electrical assessment of the action of drugs, and that a comprehensive evaluation (voltage and calcium) provides important insights of how a compound may affect cardiac function. In appreciation that some compounds may uniquely affect wave propagation and conduction velocity, we also illustrated that optogenetic point stimulation can be achieved in the 96-well plate format that may be useful in such cases, Figure 7. Current classic cardiotoxicity testing derives from assessment of torsadogenic risk of arrhythmias, which in turn is linked to QT-interval prolongation and APD prolongation. Torsade de pointes type of arrhythmias constitute a small fraction of possible arrhythmia mechanisms. The efforts to re-vamp cardiotoxicity testing in the CiPA initiative (comprehensive in-vitro proarrhythmia assay)[57–60] are motivated by desire to expand the assessment, reduce the false positives and false negatives and to arrive at better predictors of risk. HT high-content measurements provided by OptoDyCE-plate can contribute to such efforts.

The desire for higher throughput and the desire for more realistic representation of cardiac tissue structure, such as 3D oriented cell growth, are often at odds with each other. It is important to systematically validate the minimal requirements for representing cardiac syncytium *in vitro* without compromising predictions for the action of drugs. Our results with OptoDyCE using 96-well and 384-well format, monolayers grown on glass-bottom surface, as well as quisi-3D isotropic and anisotropic cell growth on electrospun matrices within such HT plates provide evidence that simple densely-grown monolayers may be a “good enough” representation for assessment of electrical responses of cardiac tissue to drugs, as qualitatively all cases agreed in the predictions. Quasi-3D growth yielded similar baseline parameters to the cell monolayers (Figures 5-6 compared to Figure 4). Structured anisotropic cardiac cell growth had slightly prolonged APD and CTD compared to isotropic growth on electrospun matrices, consistent with hypertrophic growth[35, 36], Figures 5-6. We also saw steeper restitution and tighter spread of the responses to drugs in these anisotropic constructs. Such structured cell growth may be particularly important in accurately capturing the effects of compounds on conduction properties and on cardiac mechanics, where anisotropy uniquely impacts function.

Overall, the combination of increased throughput/parallelism and enhanced information content, as demonstrated here, can help speed up testing of drugs in a more rigorous way and accelerate the development of new therapeutics. The combination of OptoDyCE-plate with robotic liquid handling and further automation of cell plating, optical sensor application and drug application can reduce sample variability and further improve the preclinical assessment of the action of genetic or pharmacological modifications.

## Acknowledgements

This work was supported in part by grants from the National Science Foundation (EFMA1830941 and CBET1705645) and the National Institutes of Health (R01HL144157). Prof. Gil Bub (McGill) helped with Arduino coding of the temporal multiplexing.

## Author Contributions

YWH, JLH and EE designed the system and the experiments. YWH finalized the design, built and characterized the system and wrote custom scripts. JLH prepared all samples and performed molecular modifications. JLH and YWH performed all experiments and analyzed the data. EE oversaw the project, helped interpret the results, and secured funding. EE, YWH, and JLH wrote and edited the paper.

## Data Availability

All data are shown in the manuscript.

## References

[1] E. National Academies of Sciences, Medicine, New Approach Methods (NAMs) for Human Health Risk Assessment: Proceedings of a Workshop–in Brief, The National Academies Press, Washington, DC, 2022.

[2] M.C. Daley, U. Mende, B.R. Choi, P.D. McMullen, K.L.K. Coulombe, Beyond pharmaceuticals: Fit-for-purpose new approach methodologies for environmental cardiotoxicity testing, Altex 40(1) (2023) 103–116.

[3] S. Schmeisser, A. Miccoli, M. von Bergen, E. Berggren, A. Braeuning, W. Busch, C. Desaintes, A. Gourmelon, R. Grafström, J. Harrill, T. Hartung, M. Herzler, G. Kass, N. Kleinstreuer, M. Leist, M. Luijten, P. Marx-Stoelting, O. Poetz, B. van Ravenzwaay, R. Roggeband, V. Rogiers, A. Roth, P. Sanders, R.S. Thomas, A. Marie Vinggaard, M. Vinken, B. van de Water, A. Luch, T. Tralau, New approach methodologies in human regulatory toxicology - Not if, but how and when!, Environ Int 178 (2023) 108082.

[4] The Precision Toxicology initiative, Toxicol Lett 383 (2023) 33-42.

[5] T.G. Bean, V.R. Beasley, P. Berny, K.M. Eisenreich, J.E. Elliott, M.L. Eng, P.C. Fuchsman, M.S. Johnson, M.D. King, R. Mateo, C.B. Meyer, C.J. Salice, B.A. Rattner, Toxicological effects assessment for wildlife in the 21st century: Review of current methods and recommendations for a path forward, Integr Environ Assess Manag (2023).

[6] W.L. McKeithan, D.A.M. Feyen, A.A.N. Bruyneel, K.J. Okolotowicz, D.A. Ryan, K.J. Sampson, F. Potet, A. Savchenko, J. Gómez-Galeno, M. Vu, R. Serrano, A.L. George, Jr., R.S. Kass, J.R. Cashman, M. Mercola, Reengineering an Antiarrhythmic Drug Using Patient hiPSC Cardiomyocytes to Improve Therapeutic Potential and Reduce Toxicity, Cell Stem Cell 27(5) (2020) 813–821.e6.

[7] W.L. McKeithan, A. Savchenko, M.S. Yu, F. Cerignoli, A.A.N. Bruyneel, J.H. Price, A.R. Colas, E.W. Miller, J.R. Cashman, M. Mercola, An Automated Platform for Assessment of Congenital and Drug-Induced Arrhythmia with hiPSC-Derived Cardiomyocytes, Frontiers in physiology 8 (2017) 766.

[8] M. Ashraf, S. Mohanan, B.R. Sim, A. Tam, K. Rahemipour, D. Brousseau, S. Thibault, A.D. Corbett, G. Bub, Random access parallel microscopy, eLife 10 (2021).

[9] R.A.B. Burton, A. Klimas, C.M. Ambrosi, J. Tomek, A. Corbett, E. Entcheva, G. Bub, Optical control of excitation waves in cardiac tissue, Nature photonics 9(12) (2015) 813–816.

[10] V. Emiliani, E. Entcheva, R. Hedrich, P. Hegemann, K.R. Konrad, C. Lüscher, M. Mahn, Z.-H. Pan, R.R. Sims, J. Vierock, O. Yizhar, Optogenetics for light control of biological systems, Nature Reviews Methods Primers 2(1) (2022) 55.

[11] E. Entcheva, M.W. Kay, Cardiac optogenetics: a decade of enlightenment, Nature reviews. Cardiology 18(5) (2021) 349–367.

[12] A. Klimas, G. Ortiz, S.C. Boggess, E.W. Miller, E. Entcheva, Multimodal on-axis platform for all-optical electrophysiology with near-infrared probes in human stem-cell-derived cardiomyocytes, Prog Biophys Mol Biol 154 (2020) 62–70.

[13] A. Klimas, C.M. Ambrosi, J. Yu, J.C. Williams, H. Bien, E. Entcheva, OptoDyCE as an automated system for high-throughput all-optical dynamic cardiac electrophysiology, Nature communications 7 (2016) 11542.

[14] C. Nguyen, H. Upadhyay, M. Murphy, G. Borja, E.J. Rozsahegyi, A. Barnett, T. Brookings, O.B. McManus, C.A. Werley, Simultaneous voltage and calcium imaging and optogenetic stimulation with high sensitivity and a wide field of view, Biomedical optics express 10(2) (2019) 789–806.

[15] C.A. Werley, M.P. Chien, A.E. Cohen, Ultrawidefield microscope for high-speed fluorescence imaging and targeted optogenetic stimulation, Biomedical optics express 8(12) (2017) 5794–5813.

[16] D.R. Hochbaum, Y. Zhao, S.L. Farhi, N. Klapoetke, C.A. Werley, V. Kapoor, P. Zou, J.M. Kralj, D. Maclaurin, N. Smedemark-Margulies, J.L. Saulnier, G.L. Boulting, C. Straub, Y.K. Cho, M. Melkonian, G.K. Wong, D.J. Harrison, V.N. Murthy, B.L. Sabatini, E.S. Boyden, R.E. Campbell, A.E. Cohen, All-optical electrophysiology in mammalian neurons using engineered microbial rhodopsins, Nat Methods 11(8) (2014) 825–33.

[17] Y.W. Heinson, J.L. Han, E. Entcheva, Portable low-cost macroscopic mapping system for all-optical cardiac electrophysiology, J Biomed Opt 28(1) (2023) 016001.

[18] C.A. Werley, S. Boccardo, A. Rigamonti, E.M. Hansson, A.E. Cohen, Multiplexed Optical Sensors in Arrayed Islands of Cells for multimodal recordings of cellular physiology, Nature communications 11(1) (2020) 3881.

[19] G.B. Borja, H. Zhang, B.N. Harwood, J. Jacques, J. Grooms, R.O. Chantre, D. Zhang, A. Barnett, C.A. Werley, Y. Lu, S.F. Nagle, O.B. McManus, G.T. Dempsey, Highly Parallelized, Multicolor Optogenetic Recordings of Cellular Activity for Therapeutic Discovery Applications in Ion Channels and Disease-Associated Excitable Cells, Front Mol Neurosci 15 (2022) 896320.

[20] A. Allan, J. Creech, C. Hausner, P. Krajcarski, B. Gunawan, N. Poulin, P. Kozlowski, C.W. Clark, R. Dow, P. Saraithong, D.B. Mair, T. Block, A. Monteiro da Rocha, D.H. Kim, T.J. Herron, High-throughput longitudinal electrophysiology screening of mature chamber-specific hiPSC-CMs using optical mapping, iScience 26(7) (2023) 107142.

[21] M.S. Ma, S. Sundaram, L. Lou, A. Agarwal, C.S. Chen, T.G. Bifano, High throughput screening system for engineered cardiac tissues, Frontiers in Bioengineering and Biotechnology 11 (2023).

[22] C. Credi, V. Balducci, U. Munagala, C. Cianca, S. Bigiarini, A.A.F. de Vries, L.M. Loew, F.S. Pavone, E. Cerbai, L. Sartiani, L. Sacconi, Fast Optical Investigation of Cardiac Electrophysiology by Parallel Detection in Multiwell Plates, Frontiers in physiology 12 (2021) 692496.

[23] L.J. Bugaj, W.A. Lim, High-throughput multicolor optogenetics in microwell plates, Nature Protocols 14(7) (2019) 2205–2228.

[24] W. Liu, J.L. Han, J. Tomek, G. Bub, E. Entcheva, Simultaneous Widefield Voltage and Dye-Free Optical Mapping Quantifies Electromechanical Waves in Human Induced Pluripotent Stem Cell-Derived Cardiomyocytes, ACS Photonics 10(4) (2023) 1070–1083.

[25] W.S. Redfern, L. Carlsson, A.S. Davis, W.G. Lynch, I. MacKenzie, S. Palethorpe, P.K. Siegl, I. Strang, A.T. Sullivan, R. Wallis, A.J. Camm, T.G. Hammond, Relationships between preclinical cardiac electrophysiology, clinical QT interval prolongation and torsade de pointes for a broad range of drugs: evidence for a provisional safety margin in drug development, Cardiovasc Res 58(1) (2003) 32–45.

[26] S.H. Ingwersen, S. Snel, T.G. Mant, D. Edwards, Nonlinear multiple-dose pharmacokinetics of the dopamine reuptake inhibitor vanoxerine, J Pharm Sci 82(11) (1993) 1164–6.

[27] S.H. Ingwersen, T.G. Mant, J.J. Larsen, Food intake increases the relative oral bioavailability of vanoxerine, British journal of clinical pharmacology 35(3) (1993) 308–10.

[28] S.A. Cherstniakova, D. Bi, D.R. Fuller, J.Z. Mojsiak, J.M. Collins, L.R. Cantilena, Metabolism of vanoxerine, 1-[2-[bis(4-fluorophenyl)methoxy]ethyl]-4-(3-phenylpropyl)piperazine, by human cytochrome P450 enzymes, Drug Metab Dispos 29(9) (2001) 1216–20.

[29] A.E. Lacerda, Y.A. Kuryshev, G.X. Yan, A.L. Waldo, A.M. Brown, Vanoxerine: cellular mechanism of a new antiarrhythmic, J Cardiovasc Electrophysiol 21(3) (2010) 301–10.

[30] N. Matsumoto, C.M. Khrestian, K. Ryu, A.E. Lacerda, A.M. Brown, A.L. Waldo, Vanoxerine, a new drug for terminating atrial fibrillation and flutter, J Cardiovasc Electrophysiol 21(3) (2010) 311–9.

[31] I. Cakulev, A.E. Lacerda, C.M. Khrestian, K. Ryu, A.M. Brown, A.L. Waldo, Oral vanoxerine prevents reinduction of atrial tachyarrhythmias: preliminary results, J Cardiovasc Electrophysiol 22(11) (2011) 1266–73.

[32] C.A. Obejero-Paz, A. Bruening-Wright, J. Kramer, P. Hawryluk, M. Tatalovic, H.C. Dittrich, A.M. Brown, Quantitative Profiling of the Effects of Vanoxerine on Human Cardiac Ion Channels and its Application to Cardiac Risk, Scientific reports 5(1) (2015) 17623.

[33] H.C. Dittrich, G.K. Feld, T.D. Bahnson, A.J. Camm, S. Golitsyn, A. Katz, J.I. Koontz, P.R. Kowey, A.L. Waldo, A.M. Brown, COR-ART: A multicenter, randomized, double-blind, placebo-controlled dose-ranging study to evaluate single oral doses of vanoxerine for conversion of recent-onset atrial fibrillation or flutter to normal sinus rhythm, Heart Rhythm 12(6) (2015) 1105–12.

[34] J.P. Piccini, E.L. Pritchett, B.A. Davison, G. Cotter, L.E. Wiener, G. Koch, G. Feld, A. Waldo, I.C. van Gelder, A.J. Camm, P.R. Kowey, J. Iwashita, H.C. Dittrich, Randomized, double-blind, placebo-controlled study to evaluate the safety and efficacy of a single oral dose of vanoxerine for the conversion of subjects with recent onset atrial fibrillation or flutter to normal sinus rhythm: RESTORE SR, Heart Rhythm 13(9) (2016) 1777–83.

[35] C.Y. Chung, H. Bien, E.A. Sobie, V. Dasari, D. McKinnon, B. Rosati, E. Entcheva, Hypertrophic phenotype in cardiac cell assemblies solely by structural cues and ensuing self-organization, Faseb j 25(3) (2011) 851–62.

[36] C.Y. Chung, H. Bien, E. Entcheva, The role of cardiac tissue alignment in modulating electrical function, J Cardiovasc Electrophysiol 18(12) (2007) 1323–9.

[37] M.A. Mandegar, N. Huebsch, E.B. Frolov, E. Shin, A. Truong, M.P. Olvera, A.H. Chan, Y. Miyaoka, K. Holmes, C.I. Spencer, L.M. Judge, D.E. Gordon, T.V. Eskildsen, J.E. Villalta, M.A. Horlbeck, L.A. Gilbert, N.J. Krogan, S.P. Sheikh, J.S. Weissman, L.S. Qi, P.L. So, B.R. Conklin, CRISPR Interference Efficiently Induces Specific and Reversible Gene Silencing in Human iPSCs, Cell Stem Cell 18(4) (2016) 541–53.

[38] N. Alerasool, D. Segal, H. Lee, M. Taipale, An efficient KRAB domain for CRISPRi applications in human cells, Nat Methods 17(11) (2020) 1093–1096.

[39] J.L. Han, Y.W. Heinson, C.J. Chua, W. Liu, E. Entcheva, CRISPRi Gene Modulation and All-Optical Electrophysiology in Post-Differentiated Human iPSC-Cardiomyocytes, bioRxiv (2023) (2023) 2023.05.07.539756.

[40] M.P. Pressler, A. Horvath, E. Entcheva, Sex-dependent transcription of cardiac electrophysiology and links to acetylation modifiers based on the GTEx database, Front Cardiovasc Med 9 (2022) 941890.

[41] M.R. Pozo, G.W. Meredith, E. Entcheva, Human iPSC-Cardiomyocytes as an Experimental Model to Study Epigenetic Modifiers of Electrophysiology, Cells 11(2) (2022).

[42] I. Kopljar, A. De Bondt, P. Vinken, A. Teisman, B. Damiano, N. Goeminne, I. Van den Wyngaert, D.J. Gallacher, H.R. Lu, Chronic drug-induced effects on contractile motion properties and cardiac biomarkers in human induced pluripotent stem cell-derived cardiomyocytes, British journal of pharmacology 174(21) (2017) 3766–3779.

[43] I. Kopljar, D.J. Gallacher, A. De Bondt, L. Cougnaud, E. Vlaminckx, I. Van den Wyngaert, H.R. Lu, Functional and Transcriptional Characterization of Histone Deacetylase Inhibitor-Mediated Cardiac Adverse Effects in Human Induced Pluripotent Stem Cell-Derived Cardiomyocytes, Stem cells translational medicine 5(5) (2016) 602–612.

[44] B. Brundel, J. Li, D. Zhang, Role of HDACs in cardiac electropathology: Therapeutic implications for atrial fibrillation, Biochim Biophys Acta Mol Cell Res 1867(3) (2020) 118459.

[45] Z. Jia, V. Valiunas, Z. Lu, H. Bien, H. Liu, H.Z. Wang, B. Rosati, P.R. Brink, I.S. Cohen, E. Entcheva, Stimulating cardiac muscle by light: cardiac optogenetics by cell delivery, Circ Arrhythm Electrophysiol 4(5) (2011) 753–60.

[46] C.J. Chua, J.L. Han, W. Li, W. Liu, E. Entcheva, Integration of Engineered “Spark-Cell” Spheroids for Optical Pacing of Cardiac Tissue, Front Bioeng Biotechnol 9 (2021) 658594.

[47] W. Li, J.L. Han, E. Entcheva, Protein and mRNA Quantification in Small Samples of Human-Induced Pluripotent Stem Cell-Derived Cardiomyocytes in 96-Well Microplates, Methods Mol Biol 2485 (2022) 15–37.

[48] A. Obergrussberger, S. Stölzle-Feix, N. Becker, A. Brüggemann, N. Fertig, C. Möller, Novel screening techniques for ion channel targeting drugs, Channels (Austin) 9(6) (2015) 367–75.

[49] W. Li, J.L. Han, E. Entcheva, Syncytium cell growth increases Kir2.1 contribution in human iPSC-cardiomyocytes, Am J Physiol Heart Circ Physiol 319(5) (2020) H1112–h1122.

[50] C. Kane, D.T. Du, N. Hellen, C.M. Terracciano, The Fallacy of Assigning Chamber Specificity to iPSC Cardiac Myocytes from Action Potential Morphology, Biophys J 110(1) (2016) 281–3.

[51] L. Doerr, U. Thomas, D.R. Guinot, C.T. Bot, S. Stoelzle-Feix, M. Beckler, M. George, N. Fertig, New easy-to-use hybrid system for extracellular potential and impedance recordings, Journal of laboratory automation 20(2) (2015) 175–88.

[52] N. Belbachir, N. Cunningham, J.C. Wu, High-Throughput Analysis of Drug Safety Responses in Induced Pluripotent Stem Cell-Derived Cardiomyocytes Using Multielectrode Array, Methods Mol Biol 2485 (2022) 99–109.

[53] G.T. Dempsey, K.W. Chaudhary, N. Atwater, C. Nguyen, B.S. Brown, J.D. McNeish, A.E. Cohen, J.M. Kralj, Cardiotoxicity screening with simultaneous optogenetic pacing, voltage imaging and calcium imaging, J Pharmacol Toxicol Methods 81 (2016) 240–50.

[54] A. Sharma, P.W. Burridge, W.L. McKeithan, R. Serrano, P. Shukla, N. Sayed, J.M. Churko, T. Kitani, H. Wu, A. Holmstrom, E. Matsa, Y. Zhang, A. Kumar, A.C. Fan, J.C. Del Alamo, S.M. Wu, J.J. Moslehi, M. Mercola, J.C. Wu, High-throughput screening of tyrosine kinase inhibitor cardiotoxicity with human induced pluripotent stem cells, Sci Transl Med 9(377) (2017).

[55] P. Dorian, D. Newman, Rate dependence of the effect of antiarrhythmic drugs delaying cardiac repolarization: an overview, EP Europace 2(4) (2000) 277–285.

[56] F. Cerignoli, D. Charlot, R. Whittaker, R. Ingermanson, P. Gehalot, A. Savchenko, D.J. Gallacher, R. Towart, J.H. Price, P.M. McDonough, M. Mercola, High throughput measurement of Ca2+ dynamics for drug risk assessment in human stem cell-derived cardiomyocytes by kinetic image cytometry, Journal of Pharmacological and Toxicological Methods 66(3) (2012) 246–256.

[57] D.G. Strauss, K. Blinova, Clinical Trials in a Dish, Trends in pharmacological sciences 38(1) (2017) 4–7.

[58] K. Blinova, J. Stohlman, J. Vicente, D. Chan, L. Johannesen, M.P. Hortigon-Vinagre, V. Zamora, G. Smith, W.J. Crumb, L. Pang, B. Lyn-Cook, J. Ross, M. Brock, S. Chvatal, D. Millard, L. Galeotti, N. Stockbridge, D.G. Strauss, Comprehensive Translational Assessment of Human-Induced Pluripotent Stem Cell Derived Cardiomyocytes for Evaluating Drug-Induced Arrhythmias, Toxicological sciences: an official journal of the Society of Toxicology 155(1) (2017) 234–247.

[59] G. Gintant, P. Burridge, L. Gepstein, S. Harding, T. Herron, C. Hong, J. Jalife, J.C. Wu, Use of Human Induced Pluripotent Stem Cell-Derived Cardiomyocytes in Preclinical Cancer Drug Cardiotoxicity Testing: A Scientific Statement From the American Heart Association, Circ Res 125(10) (2019) e75–e92.

[60] T. Colatsky, B. Fermini, G. Gintant, J.B. Pierson, P. Sager, Y. Sekino, D.G. Strauss, N. Stockbridge, The Comprehensive in Vitro Proarrhythmia Assay (CiPA) initiative - Update on progress, J Pharmacol Toxicol Methods 81 (2016) 15–20.

